# Global scale phylogeography of functional traits and microdiversity in *Prochlorococcus*

**DOI:** 10.1101/2023.01.24.525399

**Authors:** Lucas J. Ustick, Alyse A. Larkin, Adam C. Martiny

**Affiliations:** Department of Ecology and Evolutionary Biology, University of California Irvine, Irvine, CA 92697, USA; Department of Earth System Science, University of California Irvine, Irvine, CA 92697, USA

## Abstract

*Prochlorococcus* is the most numerically abundant photosynthetic organism in the surface ocean. The *Prochlorococcus* high-light and warm-water adapted ecotype (HLII) is comprised of extensive microdiversity, but specific functional differences between microdiverse sub-clades remain elusive. Here we characterized both functional and phylogenetic diversity within the HLII ecotype using Bio-GO-SHIP metagenomes. We found widespread variation in gene frequency connected to local environmental conditions. Metagenomically assembled marker genes and genomes revealed a globally distributed novel HLII haplotype defined by adaptation to chronically low P conditions (HLII-P). Environmental correlation analysis revealed different factors were driving gene abundances verses phylogenetic differences. An analysis of cultured HLII genomes and metagenomically assembled genomes revealed a subclade within HLII, which corresponded to the novel HLII-P haplotype. This work represents the first global assessment of the HLII ecotype’s phylogeography and corresponding functional differences. These findings together expand our understanding of how microdiversity structures functional differences and reveals the importance of nutrients as drivers of microdiversity in *Prochlorococcus*.

## Introduction

Microbial communities harbor vast phylogenetic diversity that is tightly linked to biogeographic partitioning of biomes. High resolution microbial diversity has been revealed through advances in sequencing technologies (1–5). Fine scale phylogenetic differences, termed microdiversity (greater than 97% 16S similarity), have been shown to distinguish physiologically unique populations (6,7). Moreover, these diverse microbial populations harbor distinct functional traits conserved across various phylogenetic depths (8). Microdiversity partitions important microbial traits such as antibiotic resistance, toxin production, nutrient uptake, and phage resistance (9–12). While there is a clear association between fine phylogenetic clusters and differential genome content (13), less is known about the specific functional differences associated with these microdiverse lineages. Functional differences between closely related microbes can confer significant competitive advantages and niche specialization (12). Thus, to better understand the eco-evolutionary processes that drive differentiation between closely related microbial populations, it is critical that we link phylogenetic microdiversity with specific functional adaptations.

In the numerically dominant and well-studied phytoplankton *Prochlorococcus*, we observe a clear phylogenetic organization of traits (14,15). *Prochlorococcus* phylogeny first diverges at the deepest taxonomic resolution by differences in adaptation to light, with a low light (LL) and high light (HL) clade (6). Within the high light clade, *Prochlorococcus* demonstrates further partitioning with a split between a high (HLII) and low temperature ecotype (HLI) (16,17). Within these well-established ecotypes, we observe phylogenetic microdiversity that follows clear spatial differentiation (18–20). This microdiversity has been linked to differences in genome content, but the specific functional differences are unknown (21). Niche differences between microdiverse subclades have largely been explained by correlative environmental relationships (22,23). It has been hypothesized that differences in adaptation to nutrient limitation (specifically to phosphorus and nitrogen limitation) may be associated with microdiverse lineages, but no clear connection between phylogenetic sub clades and functional gene content have been identified in *Prochlorococcus* (24–26).

The *Prochlorococcus* ecotype HLII is globally distributed and highly abundant in warm oligotrophic regions, making it a good model for studying the organization of microdiversity. Adaptations to low nutrient conditions vary spatially and are indicative of local nutrient conditions (27), but the presence/absence of these genes lack phylogenetic organization in our current ecotype groupings (24–26). While specifically exploring the phylogenetic distribution of *narB* (nitrate reductase) and *nirA* (nitrite reductase), the presence/absence of these genes did not display a clear evolutionary structure and had a sporadic distribution (24). It was suggested that this structure arose through vertical evolution of the traits from basal lineages followed by gene loss events. An alternate hypothesis is that nutrient acquisition traits are shared through horizontal gene transfer and thus do not follow a clear phylogenetic pattern (25,28). It is thus unclear whether nutrient adaptation is phylogenetically conserved at the microdiverse phylogenetic level, and how adaptation to variable nutrient conditions influences the biogeographic distribution of these populations.

Here, we aim to link the global microdiversity and phylogeography of *Prochlorococcus* with population-specific functional diversity. Specifically, we isolated reads that mapped to a well-studied phylogenetic marker gene to capture the phylogeography of *Prochlorococcus* and simultaneously quantified differences in the functional *Prochlorococcus* gene content from surface metagenomes. We then generated metagenomically assembled genomes (MAG) of the novel haplotypes identified. We hypothesize that functional differences within *Prochlorococcus* HLII is primarily driven by adaptation to different nutrient regimes. We present a hierarchical set of hypotheses regarding the phylogenetic structure of *Prochlorococcus* HLII microdiversity. The hypotheses are based on two different factors: how do nutrient conditions (primary nutrient limitation vs. co-limitation) and dispersal limitation (dispersal limited vs. effectively globally distributed) drive microdiversity. The first hypothesis predicts microdiversity to be driven by differences in adaptation to the primary limiting nutrient (Figure 1A). In this case, we would expect globally dispersed haplotypes that are differentiated based on single nutrient conditions, i.e. nitrogen (N) limitation vs. phosphorus (P) limitation. Our second hypothesis predicts microdiversity will be driven by combinatorial differences in the type of nutrient limitation (Figure 1B). Like the previous hypothesis, we expect unique haplotypes for each type (N, P) of primary nutrient limitation but also expect unique haplotypes for different combinations of co-limitation such as N/P limitation vs. N/Fe limitation. The third hypothesis predicts microdiversity will be driven by the primary limiting nutrient and dispersal-limited genetic drift (Figure 1C). In this case proximity of samples and the primary limiting nutrient would be the strongest drivers of microdiversity. For example, we would expect P limited populations in the North Atlantic Ocean to be phylogenetically distinct from P limited populations in other oceans. The final hypothesis predict microdiversity will be driven by both limiting nutrient type, nutrient co-limitation, and will be dispersal-limited (Figure 1D). In this case we would still expect regionally bounded populations, but both the primary limiting nutrient and co-limiting nutrients would lead to distinct populations.

**Figure 1:**
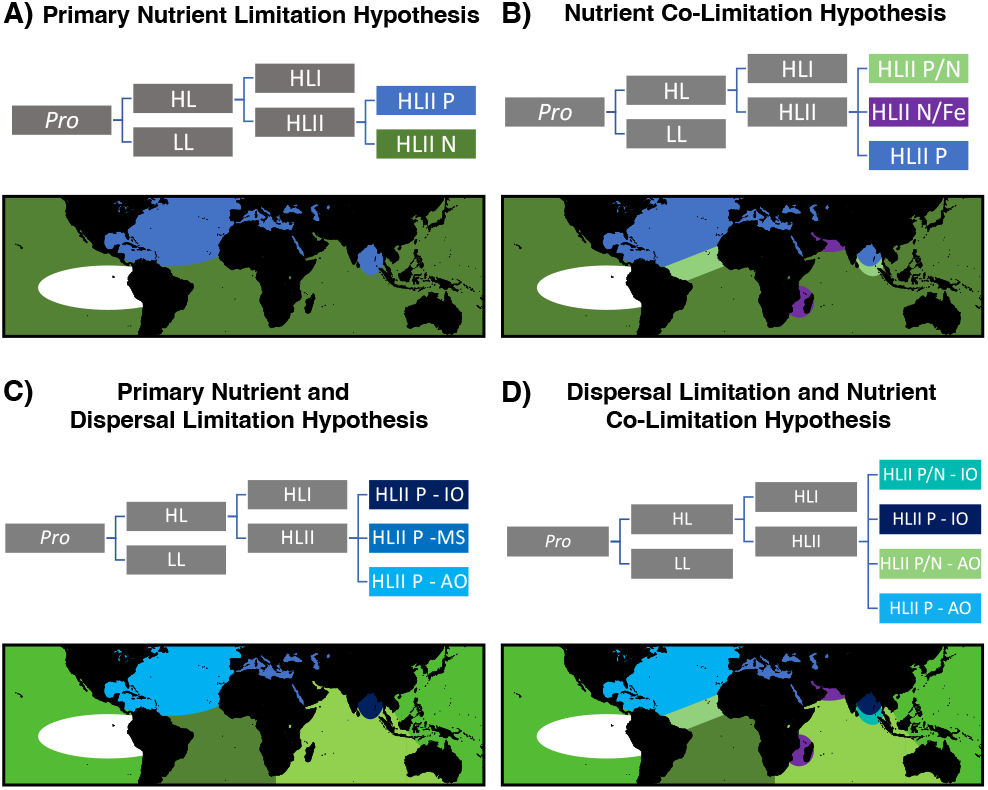
Hypotheses on the drivers of microdiversity and predicted global distributions. Hypotheses vary based on two different factors: nutrient conditions (single nutrient limitation vs. co-limitation) and dispersal limitation (dispersal limited vs. effectively globally distributed). Colors on the trees and maps correspond across all panels and represent the expected phylogenetic structure and distributions based on each hypothesis. (A) Primary nutrient limitation hypothesis. (B) Nutrient co-limitation hypothesis. (C) Primary nutrient and dispersal limitation hypothesis. (D) Dispersal limitation and nutrient co-limitation hypothesis.

## Results

We analyzed 630 surface ocean metagenomes from Bio-GO-SHIP (29), and GEOTRACES (30) and isolated reads that mapped to *Prochlorococcus* HLII reference genomes to link phylogenetic and functional diversity at the microdiverse population level (Figure S1). We first characterized the global functional diversity within HLII by quantifying the variation in gene frequency across the global ocean, which we will refer to as genomic diversity in this study. We then related this functional diversity to phylogenetic changes by identifying dominant single nucleotide polymorphisms (SNPs) among *Prochlorococcus* populations, which we refer to as phylogenetic diversity. Based on the relationship of genomic and phylogenetic diversity, we then created targeted metagenomically assembled genomes to better understand the genomic structure of each distinct cluster.

*Prochlorococcus* genomic functional diversity revealed clear separation between major ocean regions. We performed a PCA analysis on the normalized abundance of the variable gene content within the HLII ecotype. To contextualize this dimensional reduction, we fit the PCA with temperature and overlayed the average direction and magnitude of nutrient acquisition and metabolism genes grouped by type (Fe/P/N) (Figure 2A,2B). The top 100 genes that contributed to each principal component were identified and counted based on NCBI clusters of orthologous groups annotations (COGs). PC1 captured 13% of the total variance in gene content and had the strongest correlation with temperature (Spearman rho=0.74, p<0.05)(Figure 2A, S2A, Table S1). PC1 was primarily informed by genes annotated as H (coenzyme metabolism), J (translation), and L (replication recombination and repair)(Figure 2C). Three of the top five genes when ordered by PC1 loadings were tRNA synthetases (Cysteinyl-tRNA synthetase, Histidyl-tRNA synthetase, and Seryl-tRNA synthetase). This along with the correlation with temperature suggested these represented differences in energetic adaptations. PC2 captured 8% of the total variance and was enriched in the Atlantic but depleted in the Indian Ocean Samples (Figure 2A, S2B, Table S1). PC2 was primarily informed by genes annotated as E (amino acid metabolism and transport), G (carbohydrate transport and metabolism), and H (coenzyme metabolism). PC3 captured 6% of the total variance and linked to genes annotated as M (cell wall/ membrane/ envelop biogenesis)(Figure 2, S2C). PC4 captured 4% of the total variance and was mainly informed by genes annotated as P (inorganic ion transport and metabolism). Of the top 50 genes ranked by PC4 loadings, 10 were involved in phosphorus acquisition and metabolism with positive loadings, and 4 were iron acquisition genes with negative loadings (Table S2). This relationship was further highlighted as the frequency of all acquisition and metabolism genes for phosphorus also had a strong positive relationship with PC4 (Spearman rho=0.88, p<0.05) and the average abundance of all iron genes had a negative relationship (Spearman rho=-0.79, p<0.05) (Figure 2B, S2D, Table S1). Overall, functional gene content was correlated with differences in temperature and nutrient regimes leading to distinct ocean basin distributions.

**Figure 2:**
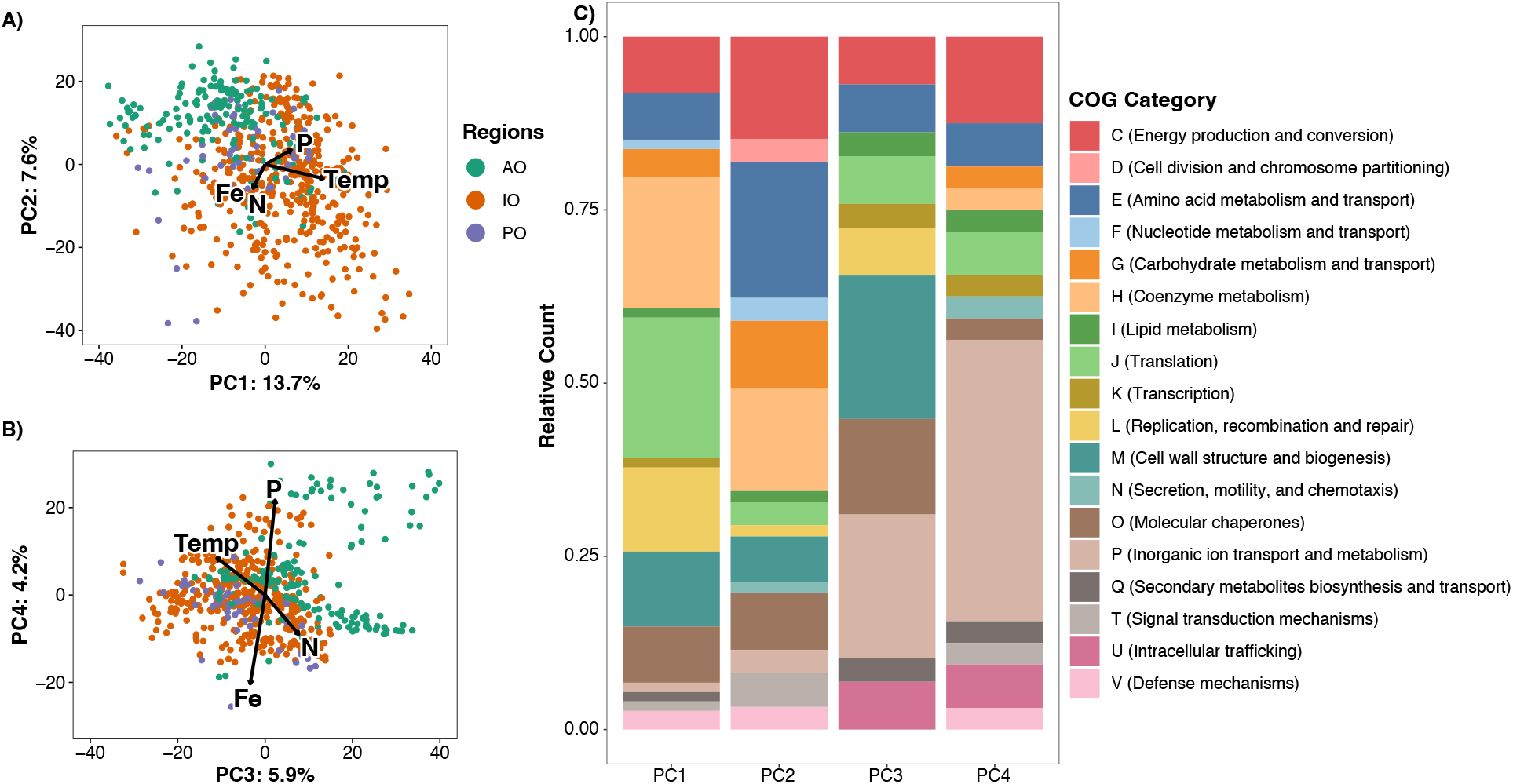
Global Variation in Gene Content (Genomic Diversity) of *Prochlorococcus* HLII. (A,B) Principal component analysis of HLII gene abundance (genomic diversity). Vector length of temperature is based on variance explained by the environmental factor, while Fe, P, and N is the average vector of nutrient acquisition genes grouped by type. Samples are colored by ocean basin Atlantic Ocean (AO), Indian Ocean (IO), and the Pacific Ocean (PO). (C) 100 most informative genes for each principal component based on PCA loadings grouped by cluster of orthologous groups (COG) annotations.

Phylogenetic microdiversity of *Prochlorococcus* populations also showed systematic biogeographic distributions. We identified 114 SNPs in the marker gene *rpoC1* globally (Figure 3A). Based on the presence and absence of these SNPs, we detected two stable clusters with strong support (63% and 92% bootstraps) and two unstable clusters (51% and 10% of bootstraps) (Figure 3A). The majority cluster was globally ubiquitous (cluster 1, present in 63% of bootstraps). A second highly conserved cluster (present in 92% bootstraps and 14 unique SNPs) was detected primarily in the North Atlantic Ocean (lat: 16-39° N, lon: 52-71° W) and intermittently in NE Indian Ocean (lat: 9-13° N lon: 85-89° E) (Figure 3A, 3B). This cluster also appeared sporadically in the central and SW Atlantic Ocean. We compared the average relative abundance of nutrient stress genes grouped based on nutrient type (P, N, Fe), termed Ω (27), to the stable SNP clusters. We labeled the second cluster HLII-P because it was associated with significantly higher abundance of phosphorus genes than cluster 1 (ΩP mean HLII-P = 1.5, Cluster 1 = −0.5, p < 0.001) as well as significantly lower abundance of iron genes (ΩFe mean HLII-P = −0.6, Cluster 1 = −0.03, t-test p < 0.001)(Figure 3A, S3). However, only 14% of functional genomic variation within HLII can be explained by phylogeny (R_mantel_ = 0.38, p < 0.001). Cluster HLII-P had a significantly higher functional genome PC4 values than cluster 1 (Cluster 1 PC4 mean = −1.60, HLII-P PC4 mean = 20.79, t-test p < 0.001), suggesting that HLII-P spatially co-occurred wherever PC4 was enriched (Figure 3A, 3B, S2, S3). Thus, we hypothesized that the functional genome PC4 represented the phylogenetically conserved functional type of HLII-P. Thus, the phylogeography of our samples indicated a strong link between phylogeny and phosphorus limitation.

**Figure 3:**
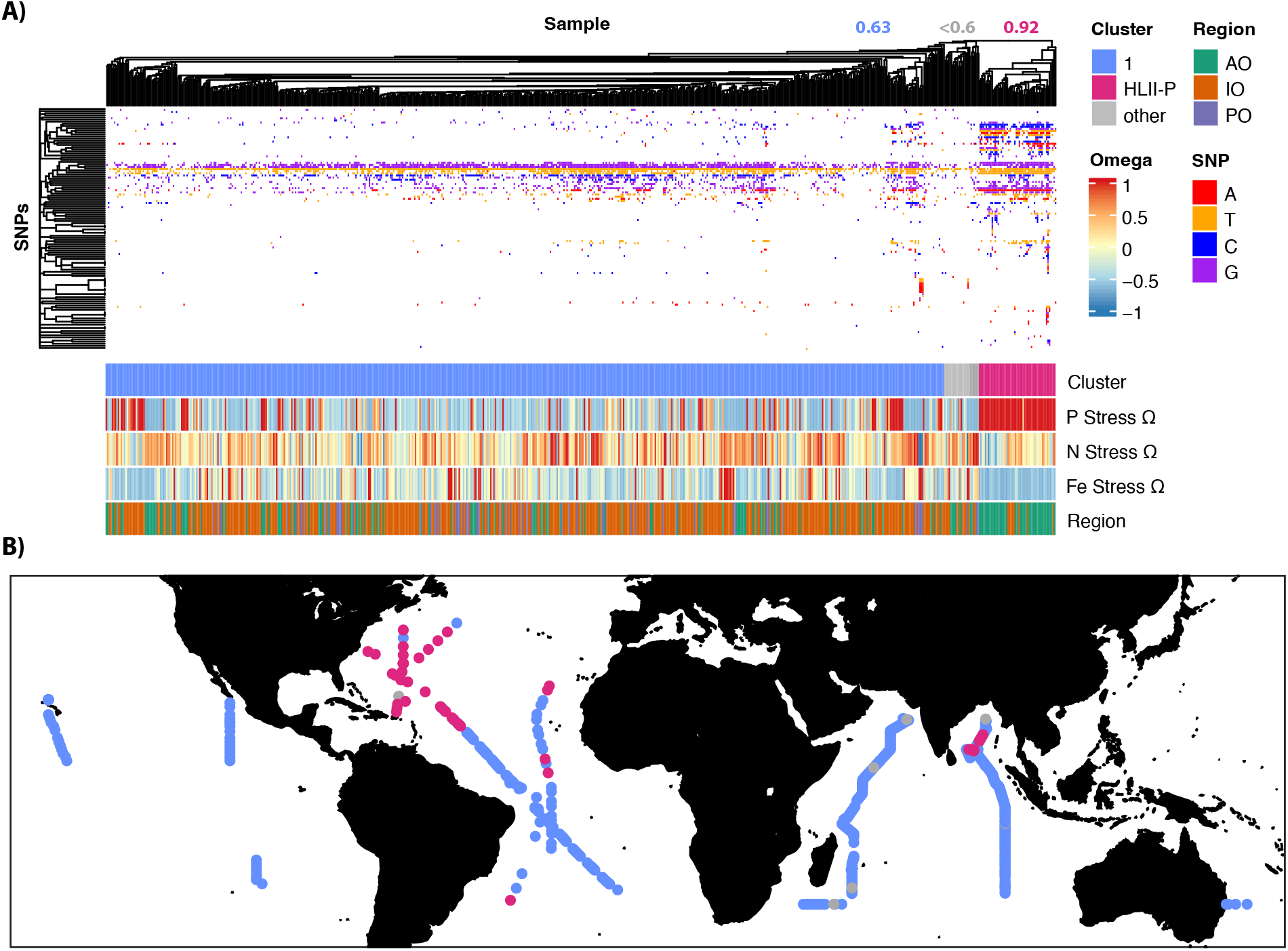
Global *rpoC1* derived Phylogenetic Diversity of *Prochlorococcus* HLII. (A) Clustering of the *rpoC1* marker gene based on sequence similarity, with corresponding region and metagenomically derived nutrient stress (Ω). Columns represent different single nucleotide polymorphisms (SNPs) with rows showing different metagenomic samples. (B) Spatial distribution of groups based on the clustering of *rpoC1* consensus sequences.

A comparison between our metagenomically derived haplotypes and amplicon sequences revealed consistency between the two assessments. We evaluated our metagenomically derived phylogenetic clusters against previously characterized amplicon derived haplotype abundances in the Indian Ocean (31). These Indan Ocean haplotypes were derived based on the relative abundance of mutually exclusive SNPs in amplicon sequences of the full *rpoC1* gene. This comparison was made to both validate the metagenomic results and to try an identify whether the HLII-P cluster was a result of one *Prochlorococcus* population or multiple co-existing populations. The HLII-P cluster was closely associated with the amplicon derived IO HLII.2 haplotype and had significantly higher relative abundances of the haplotype compared to samples in Cluster 1 (mean relative IO HLII.2 abundance, HLII-P cluster = 0.58, Cluster 1 = 0.36)(Figure S4). All samples in the HLII-P cluster had over 50% relative abundance of the amplicon derived IO HLII.2 haplotype. Differences in gene content (Genomic Diversity PC4) also had a significant linear correlation with the relative abundance of IO HLII.2 (Pearson R = 0.53, p < 0.001)(Figure S4). This together suggests the metagenomically derived HLII-P cluster is a result of a single *Prochlorococcus* population and further connects it to P limitation.

Global shifts in functional and phylogenetic diversity within *Prochlorococcus* HLII were explained by unique environmental factors. We calculated the variance in functional diversity and phylogenetic diversity explained by a variety of independent variables (Figure 4A). In our PERMANOVA analysis, we first captured the variance explained by physical distance to remove any distance effects. Then we ordered the remaining variables based on the variance explained, ordering from most to least variance explained. 4.2% of functional diversity and 1.9% of phylogenetic diversity could be explained by spatial distance between samples (Figure 4A). Once distance effects were accounted for, temperature explained the most variance in genomic diversity (3.2%), while P gene abundance (Ω P) explained the most phylogenetic diversity (2.5%) (Figure 4A). Overall, we could explain more of the genomic diversity (15%) with our environmental factors than phylogenetic diversity (8%). We also observed a stronger distance decay relationship between samples based on genomic distance (slope = 6.6*10^-4^) than phylogenetic distance (slope = 3.7*10^-7^) (Figure 4B, 4C). Overall, we observed different factors correlated with genomic differences versus phylogenetic differences, with genomic distance following regional differences such as temperature gradients while phylogenetic variation was clearly linked to phosphorus limitation.

**Figure 4:**
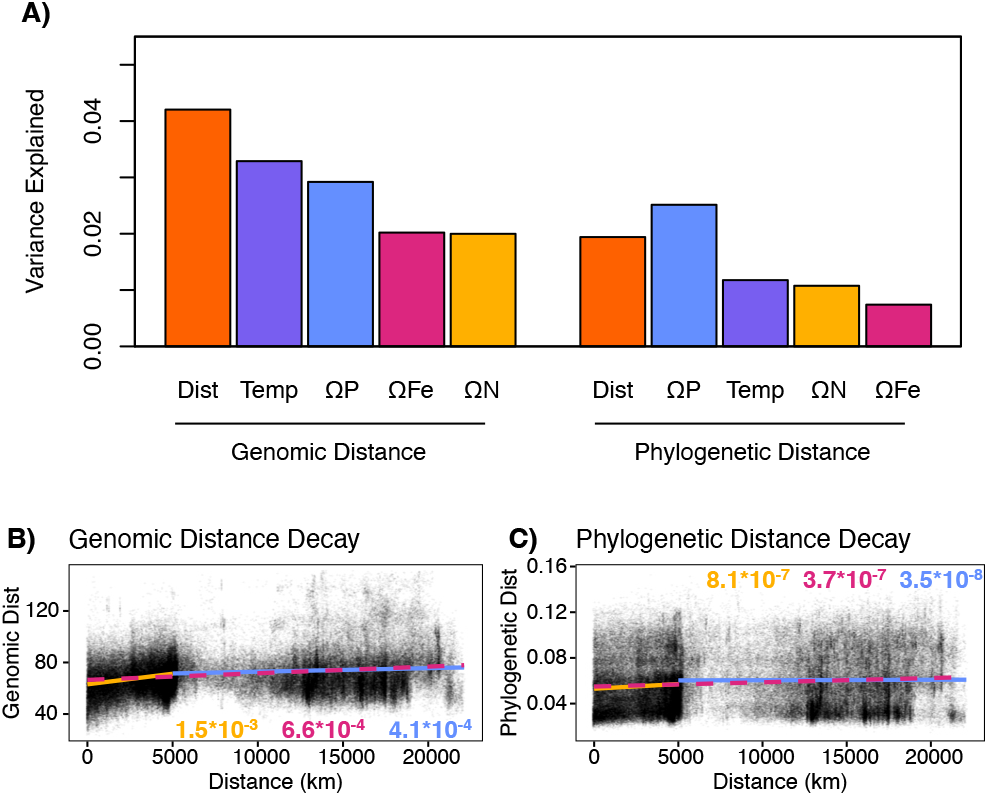
Variance explained by environmental factors and distance model. (A) Variance explained by environmental factors based on PERMANOVA. Variables are presented in the same order as they were used in the model. All relationships shown are significant (p-value < 0.01). Distance, temperature, P-stress (ΩP), N-stress (ΩN), and Fe-stress (ΩFe) were included in the PERMANOVA. (B,C) Distance decay of genomic distance (B) and phylogenetic distance (C). Sample points colored based on scatterplot density. Overall linear fit (red and dashed), within sampling transect linear fit (yellow), and between sampling transects linear fit (blue). Slopes of fits shown within figure.

Metagenomically assembled genomes (MAG) supported that HLII-P represented a phylotype adapted to low phosphorus conditions. We assembled MAGs from samples enriched with the HLII-P cluster to better understand the phylogenetic conservation of the P stress microdiverse lineage of *Prochlorococcus*. Because of the great diversity of *Prochlorococcus*, it is notoriously difficult to assemble, but we present the best assemblies which contained P acquisitions genes. Each MAG revealed an ordering of P acquisition genes consistent with other HLII reference genomes (Figure 5A). Additionally, all MAGs contained the *phoA* gene that was otherwise only found in *Prochlorococcus* HLII cultures isolated from chronically P limited regions (MIT9314, MIT9312, RS50). The parallel genomic structure of phosphorus genes in our MAGs and the reference genomes suggested a common origin of the genes. We compared the presence/absence of the phosphorus acquisition genes found in the MAG IO9-1 with cultured *Prochorococcus* HLII genomes (Figure 5B). We observed a hierarchy of presence/absence of these genes and grouped the genomes based on this pattern (Figure 5B). IO9-1 was the only MAG included in this analysis because it was the only MAG with a fully assembled *rpoC1* marker gene. A phylogenetic analysis based on the *rpoC1* gene reveals a clade that contains genomes with the full or majority P acquisition gene set including MIT9314, RS50, and the IO9-1 MAG from the HLII-P cluster (Figure 5C). The only exception was MIT9312, but this strain was isolated at 135 m depth. Thus, the outlier position may be due to an additional phylogenetic structuring of light acquisition within HLII. Our analysis of our HLII-P MAGs revealed a sub-ecotype organization of P acquisition genes that corresponded to a clade within our cultured genomes.

**Figure 5:**
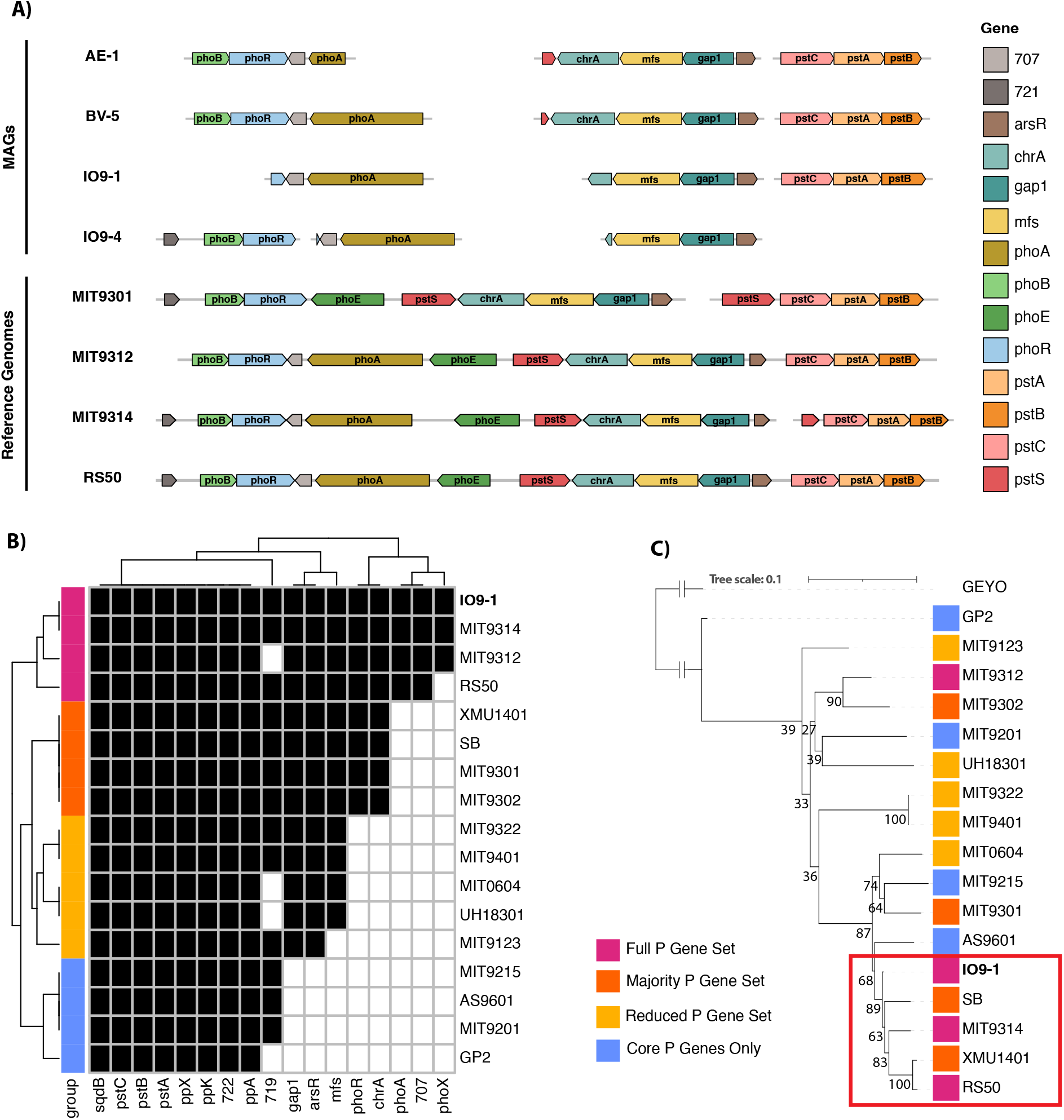
*Prochlorococcus* HLII-P MAG annotations of P acquisition genes and comparison to cultured genomes. (A) P acquisition gene presence and organization in MAG’s and cultured genomes. (B) Clustering of genomes based on presence/absence of P acquisition genes present in the MAG IO9-1. (C) Phylogenetic tree based on the *rpoC1* gene with the *Synechococcus* genome GEYO as an outlier. Bootstraps shown at the corresponding node.

## Discussion

We observed widespread functional genome diversity variation with a partial link to phylogeny. As previous studies observed a connection between microdiversity in *Prochlorococcus* and nutrient limitation, we hypothesized within ecotype functional differences would mirror these patterns and primarily be driven by differences in nutrient conditions (15,31). In our analysis, temperature explained the most variation in genome content within the ecotype HLII, but adaptation to P limitation had the strongest link to phylogeny (Figure 4). We hypothesized that differences in genomic diversity would be primarily explained by nutrient conditions. However, this hypothesis deserves revision given overall genome content was better explained by temperature (Figure 1, 4). This result was unexpected because temperature adaptation differentiates the HLII clade (high temperature) from the HLI clade (low temperature), but consistent with previous amplicon-based analyses that identified microdiverse clades within HLII associated with variable temperature niches (19). The importance of tRNA synthetase genes and temperature in the gene analysis suggests these differences are linked to energetic requirements (32). (Figure 2, 4). Previous work has linked phylogenetic marker genes to environmental processes that may have overestimated the total effect of nutrient conditions on *Prochlorococcus* genomic diversity (15). This highlights the importance of evaluating both genomic changes along with phylogenetic changes because functional traits may not follow phylogeny due to processes such as horizontal gene transfer and loss of traits through genomic streamlining (33).

Our analysis supports the existence of a novel HLII P-stress haplotype with implications for the mechanism and evolutionary history of microdiversity within *Prochlorococcus*. The organization of phosphorus uptake genes does not follow phylogeny at the broader ecotype level in *Prochlorococcus* and other marine lineages such as *Roseobacter* (25,34). This pattern is common across different microbes and supports the hypothesis that traits organize phylogenetically based on a hierarchy of biochemical complexity, with a “simple” trait like P acquisition being conserved only at the microdiverse level (8,15,35). Our analysis reveals a clear organization of P acquisition genes within the HLII ecotype highlighting the importance of phylogenetic assessment across phylogenetic depths (Figure 3, 5, S3, S4). This contrasts with nitrogen acquisition genes which have not shown any clear within ecotype organization (24,26). For nitrogen acquisition genes, it has been hypothesized that they evolved vertically then were lost or selected for in more recent lineages causing the sporadic distribution of the trait in *Prochlorococcus* ecotypes (24). While P acquisition genes have been found in *Prochlorococcus* phage genomes (28), our analysis suggests the full set of P acquisition genes is phylogenetically conserved in locations of extreme P limitation. Gene abundance of N acquisition genes is variable and changes along a continuous gradient globally, while the presence of P acquisition genes are spatially confined to a few distinct regions (27). The difference in distributions could suggest different functional and eco-evolutionary processes are acting on P acquisition versus N acquisition genes. Novel microdiverse sub-taxa can evolve by either the acquisition of a new trait or shifting growth optima along a single trait axis (12). This might also explain why only 14% of genomic changes could be explained by phylogeny. The rest of the variation may be due to regional differences in N limitation, while P limited regions appear to be stable. These results support our primary nutrient limitation hypothesis, which predicted microdiversity will be driven by differences in adaptation to the primary limiting nutrient and the resulting haplotypes will be globally dispersed (Figure 1A).

When comparing the amount of variance explained by spatial autocorrelation, we observed a stronger distance decay relationship between gene content than phylogenetic differences (Figure 4B, 4C). Microdiverse differences in phylogeny may not show spatial-auto correlation because dispersal overcomes drift in a manner similar to the Bass-Becking hypothesis, ‘everything is everywhere but the environment selects’ (36,37). The pattern we observed is indicative of strong contemporary effects combining selection and rapid dispersal resulting in globally conserved microdiverse haplotypes (38). Alternatively, our metagenomically derived phylogeny may not have the resolution to capture drift between these populations resulting in spatial autocorrelation masked by noisy data.

While metagenomics does allow for many comprehensive analyses, there are also various caveats related to these analyses. To overcome sequencing error and short read length we used a mapping-based consensus method. This method cannot capture underlying diversity in non-dominant sequence types (39). *Prochlorococcus* has been shown to have multiple co-existing microdiverse haplotypes *in situ*, which vary in abundance (31). Despite these caveats, our analysis of amplicon data in the Indian Ocean suggest the HLII-P haplotype is a single haplotype and the phylogenetic signal is not due to multiple co-existing haplotypes (Figure S4). There are also some caveats when interpreting MAGs. Due to the fact that *Prochlorococcus* populations are often made up of multiple closely related haplotypes, it is difficult to create complete assemblies that find a consistent path through the assembly graph, resulting in fragmentation (40). While we postulate that the consistent assembly structure of P acquisition genes in our MAGs is a sign of selection, it could also be a result of divergence in unassembled regions that are not captured in our assemblies.

Connecting microdiversity with specific functional groups has been limited in the field of microbial ecology due to the reliance on amplicon sequencing (41). Here, we hypothesized that differences in gene content would be primarily driven by nutrient conditions and proposed four hypotheses on the selective pressures that drive *Prochlorococcus* microdiversity. While differences in genome content were better explained by regional differences such as temperature, phylogenetic microdiversity was clearly linked to nutrient conditions with globally conserved haplotypes. This work is an example of how we can leverage large metagenomic datasets to capture global patterns that may otherwise be obscured in a local study. This analysis is the first direct link between phylogenetic microdiversity and functional diversity revealing a novel phylotype linked to nutrient stress. Quantifying the connection between microdiverse haplotypes and traits is important to link microbial processes with larger ecosystem ecology (42).

## Methods

### Metagenomes

We analyzed surface metagenomes (<25 m depth) from Bio-GO-SHIP (29), and GEOTRACES (30).

### Read Recruitment and Quality Filtering

Raw metagenomic reads were quality filtered and adapter sequences were removed using Trimmomatic v0.35 (43). Metagenomic reads were recruited using Bowtie2 v2.2.7 (44). The reads were recruited to a reference database comprised of 115 genomes with representatives of each major ecotype of Prochlorococcus and as well as Synechococcus, Pelagibacter and Roseobacter to help reduce miss recruitment of closely related microbes (Table S3). Bowtie2 was used with the following flags --no-unal --local -D 15 -R 2 -L 15 -N 1 --gbar 1 --mp 3. Resulting SAM files were sorted and indexed with samtools v1.3 into BAM files (45).

### Profiling Recruited Reads

Recruited reads were profiled using Anvi’o v5 (46). All open reading frames in the reference database were aligned and clustered using NCBI BLAST (47) and MCL (48) following the Anvi’o Pangenomic workflow (49). All gene clusters from the HLII genomes were extracted and separated into single copy core genes (SCCG) and genes in the flexible genome (non-SCCG).

### Metagenomic rpoC1 Consensus Sequences

Reads which recruited to the *rpoC1* gene across all reference genomes were extracted. Reads were then separated by sample and ecotype and all reads that mapped to HLII reference genomes were aligned to a reference *rpoC1* sequence. Based on the alignment we calculated a consensus *rpoC1* sequence for the HLII ecotype for each sample in our dataset. Consensus sequences were made by aligning with Bowtie2 (44), then the consensus was calculated and quality controlled by samtools (45). Only samples that passed the following quality metrics were used in all further analysis: a minimum of 2000 reads that mapped to *rpoC1*, minimum of 5x SCCG coverage of HLII, and the sample must have at least 90% of SCCG reads binned as HLII.

### Gene Analysis

Sequence coverage for all non-SCCG’s were normalized by average SCCG coverage. This roughly estimates the copies of a gene that are present per individual genome in the sample. We calculated a z-score for the normalized abundance of each gene and the resulting normalized coverage was analyzed using PCA analysis in R (50). We then extracted the top 100 genes that contributed to the first 4 principal components and annotated the NCBI COG for each gene using based on the Anvi’o profile data. In addition, we annotated the nutrient acquisition and metabolism genes shown in Table S2 and extracted the loadings of these genes from the aforementioned PCA analysis. Then the average magnitude and angle of each gene’s eigen vector was calculate based on groupings by nutrient type (P, N, Fe). This average vector was overlayed on the PCA along with the fit of temperature calculated using the envfit command from the vegan package in R.

### Metagenomic rpoCI Consensus Analysis

Consensus sequences were analyzed using R (50). Consensus sequences were transformed into binary sequences. Phylogenetic distances between samples was calculated using a binary Jaccard from the Vegan R package (51). The sequences were hierarchically clustered using hclust with the McQuitty method. The number of stable clusters was selected by minimizing the sum of squares within each cluster. Resulting clustering was then bootstrapped 1000 times and clusters that were in less than 60% of the bootstraps were removed from further analysis.

### Nutrient Stress Indicator

We used a genomic indicator of nutrient stress termed (Ω P) derived from *Prochlorococcus* populations. Macronutrients are below detection limits in much of the oligotrophic surface ocean making this indicator especially useful in these regions. The indicators have been linked to surface nutrient concentrations and have been previously described (27,52,53). We followed the same pipeline described in (27) and used alkaline phosphatase genes (*phoA*, *phoX*) for P-stress (Ω P), nitrite and nitrate assimilation and uptake genes (*focA*, *moaA-E*, *moeA*, *napA*, *narB*, *nirA*) for N-stress (Ω N), and Fe uptake transporter genes (*cirA*, *expD*, *febB*, *fepB/C*, *tolQ*, *tonB*) for Fe-stress (Ω Fe). The coverage of each gene was normalized to *Prochlorococcus* single copy core gene coverage (SCCG). We then calculated a Z-score for each gene and took the average across each nutrient type (P, N, Fe).

### Statistical Analysis

Spearman correlation between principal components and environmental factors was calculated in R. Pairwise physical distance between samples was modeled in R. The distance was calculated by transforming a global map into a raster image with 400 rows and 800 columns (~0.45 degree squares). Raster squares that fell on land were made impassible and the shortest distance was calculated between all samples in a pairwise fashion. We then took our distance matrix and decomposed this into a single continuous component using metaMDS in R. This was done so the distance effect could be included in our PERMANOVA analysis. Environmental variables were correlated to phylogenetic distance and functional distance using PERMANOVA analysis with the adonis2 package in R. Distance decay was estimated by extracting the slope of a linear model between genomic/phylogenetic distance and physical distance. This was done in R with the lm function.

### Metagenomically Assembled Genomes

Metagenomically assembled genomes (MAGs) were created using the following pipeline. The reads were quality controlled using the same method described previously and raw assemblies were made using the de novo assembler SPAdes with the default parameters (54). Samples were assembled individually using the metaSPAdes assembly pipeline. Resulting assembled contigs were binned using MetaBAT2 with the default parameters (55). MAGs were quality controlled and rapidly assessed using checkM (56). MAG annotations were made by aligning the resulting bins against a reference of phosphorus acquisition genes using BLAST (47).

### MAGs Phylogenetic Analysis

A phylogeny of the MAGs was created using the *rpoC1* gene. Sequences of the *rpoC1* gene were extracted from all *Prochlorococcus* reference genomes and a single *Synechococcus* genome as an outgroup. The sequences were aligned using Mega7 (57) and Muscle (58). The tree model was selected based on the maximum likelihood fit of 24 different nucleotide substitution models using MEGA7. GTR + G + I was selected since it had the lowest Bayesian Information Criterion and Akaike Information Criterion values. The phylogenetic tree was created using raxml (59) with the following arguments raxmlHPC -f a -x 318420 -p 318420 -N 100 -m GTRGAMMAX -O -o GEYO- Syn_CRD1_53540 -n out_file -s align_file -w out_dir. GEYO was a *Synechococcus* genome used as an outgroup. The resulting tree was visualized using iTOL (60).

## Author contributions

L.J.U., A.A.L., and A.C.M. designed the study. L.J.U. wrote the manuscript with input from all authors. L.J.U. analyzed the data. A.C.M. supervised the study.

## Data Availability Statement

All sequence data analyzed during the study are publicly accessible through NCBI. Raw metagenomic reads can be found through NCBI: Bio-GO-SHIP BioProject ID PRJNA656268 https://www.ncbi.nlm.nih.gov/bioproject?term=PRJNA656268, GEOTRACES BioProject PRJNA385854 and PRJNA385855 https://www.ncbi.nlm.nih.gov/bioproject?term=PRJNA385854.

## Supplemental Figures and Tables

**Figure S1:**
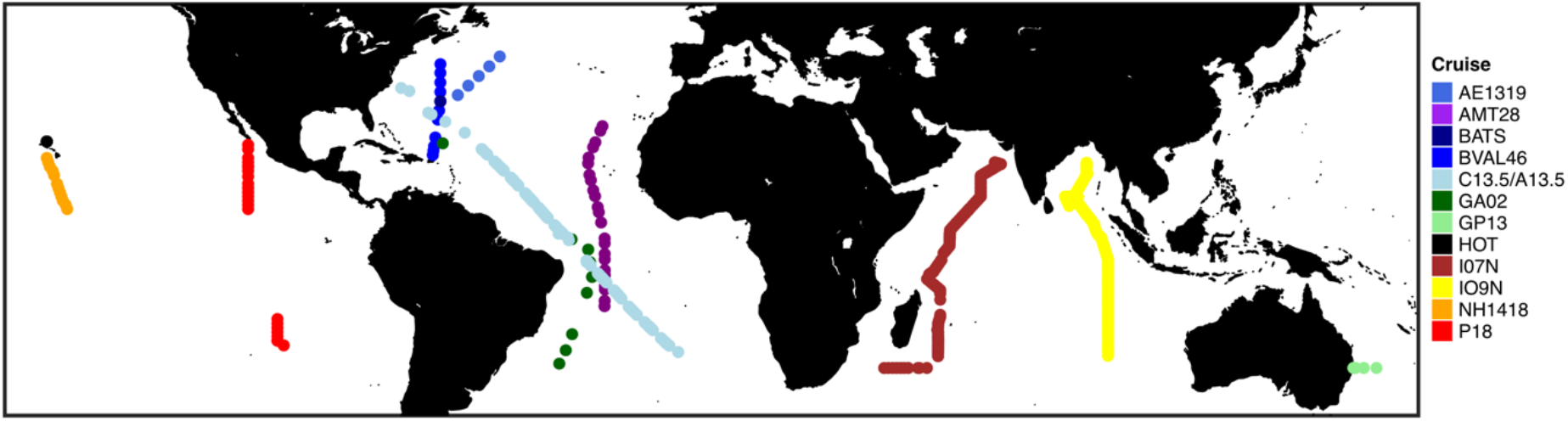
Map of metagenomic samples colored by cruise track. Bio-GO-SHIP (AE1319, AMT28, BVAL46, C13.5, IO7N, IO9N, NH1418, P18). GEOTRACES (GA02, GP13).

**Figure S2:**
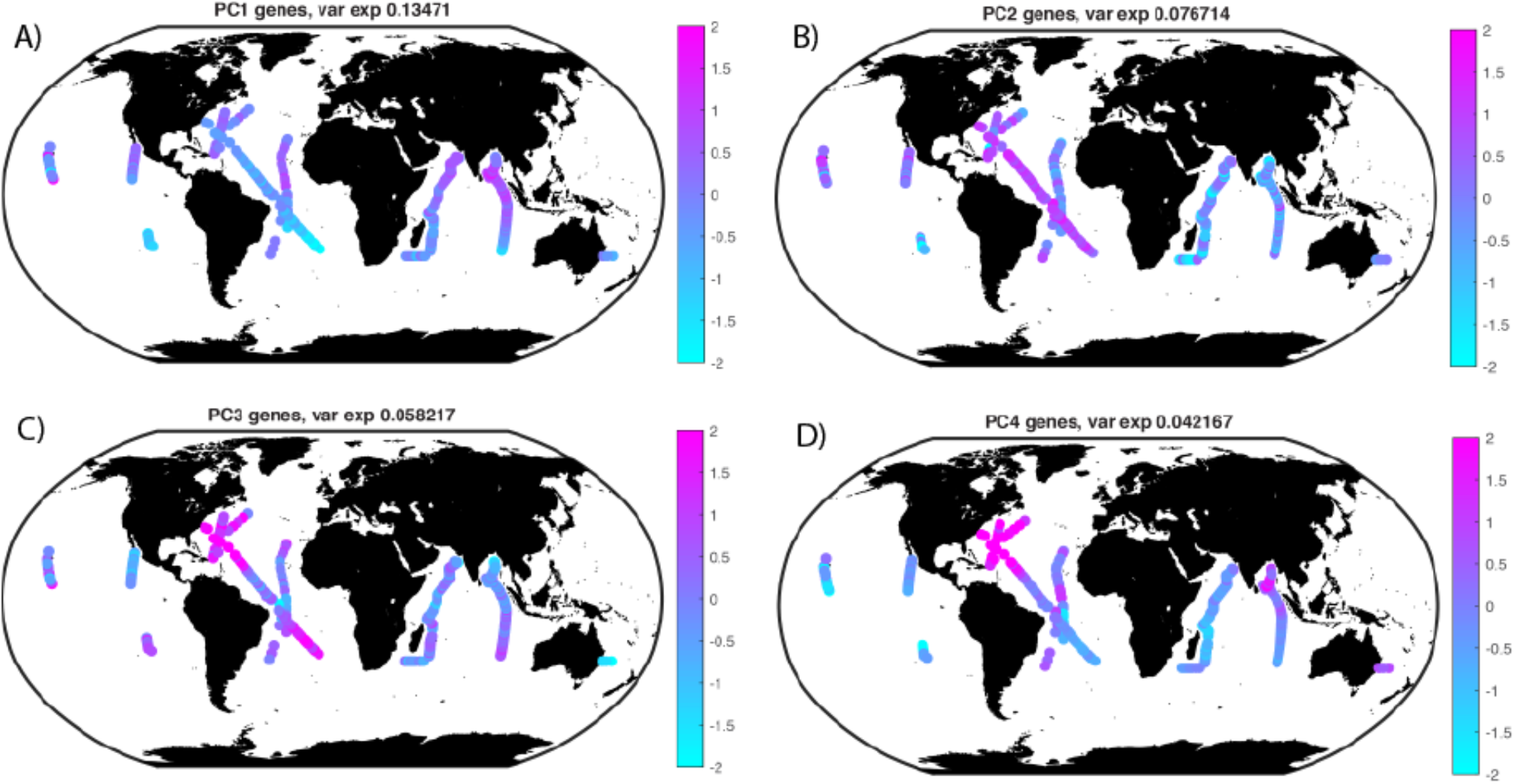
Global Variation in Gene Content (Genomic Diversity) of *Prochlorococcus* HLII. Spatial distribution of principal components based on gene abundance (Figure 2).

**Figure S3:**
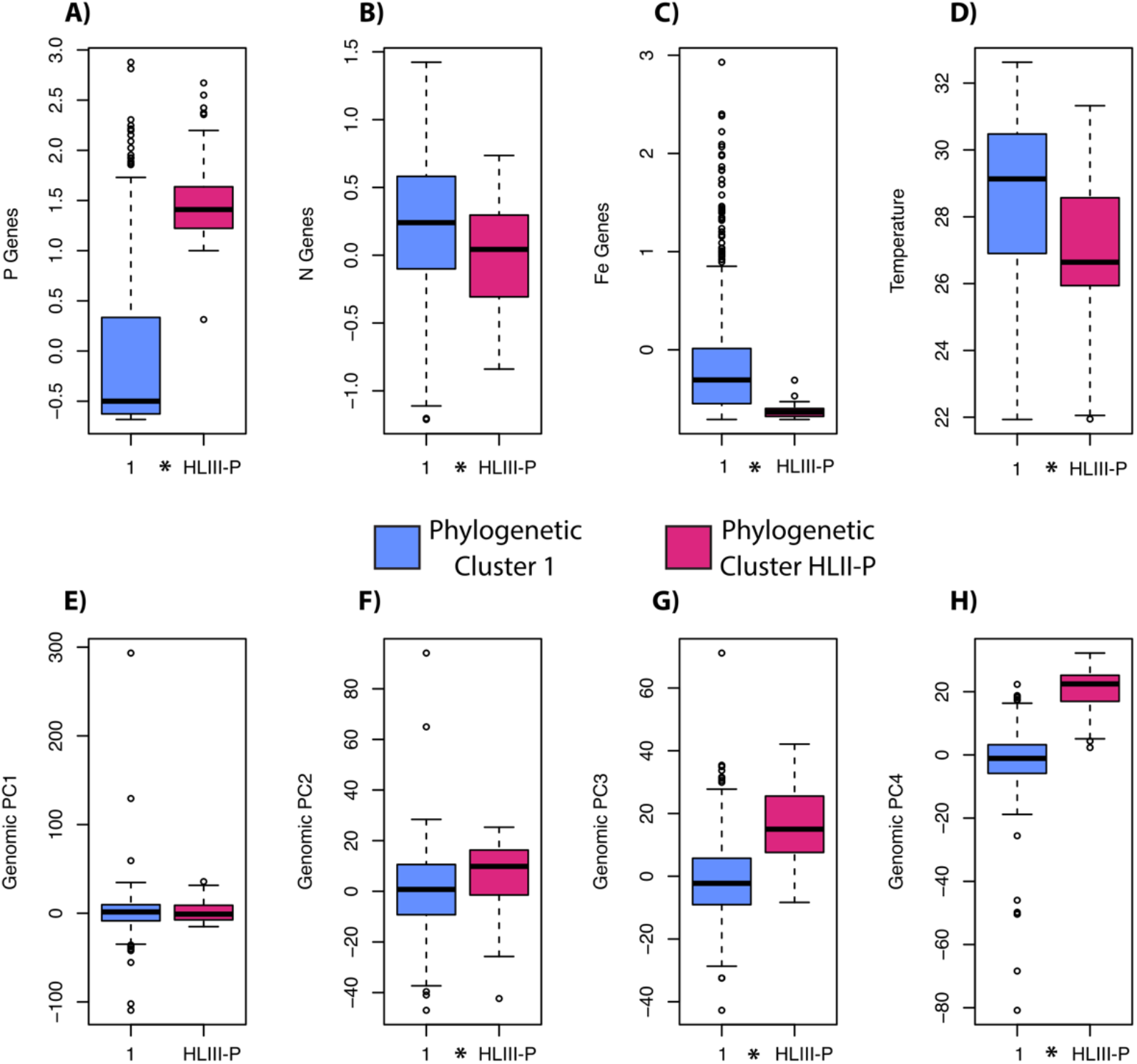
Comparison between phylogenetic clusters. (A-C) differences in average nutrient gene abundances (Table S2). (D) Average temperature between clusters. (E-H) Genomic PCA differences between phylogenetic clusters. * Denotes significant differences (T-test p value < 0.05).

**Figure S4:**
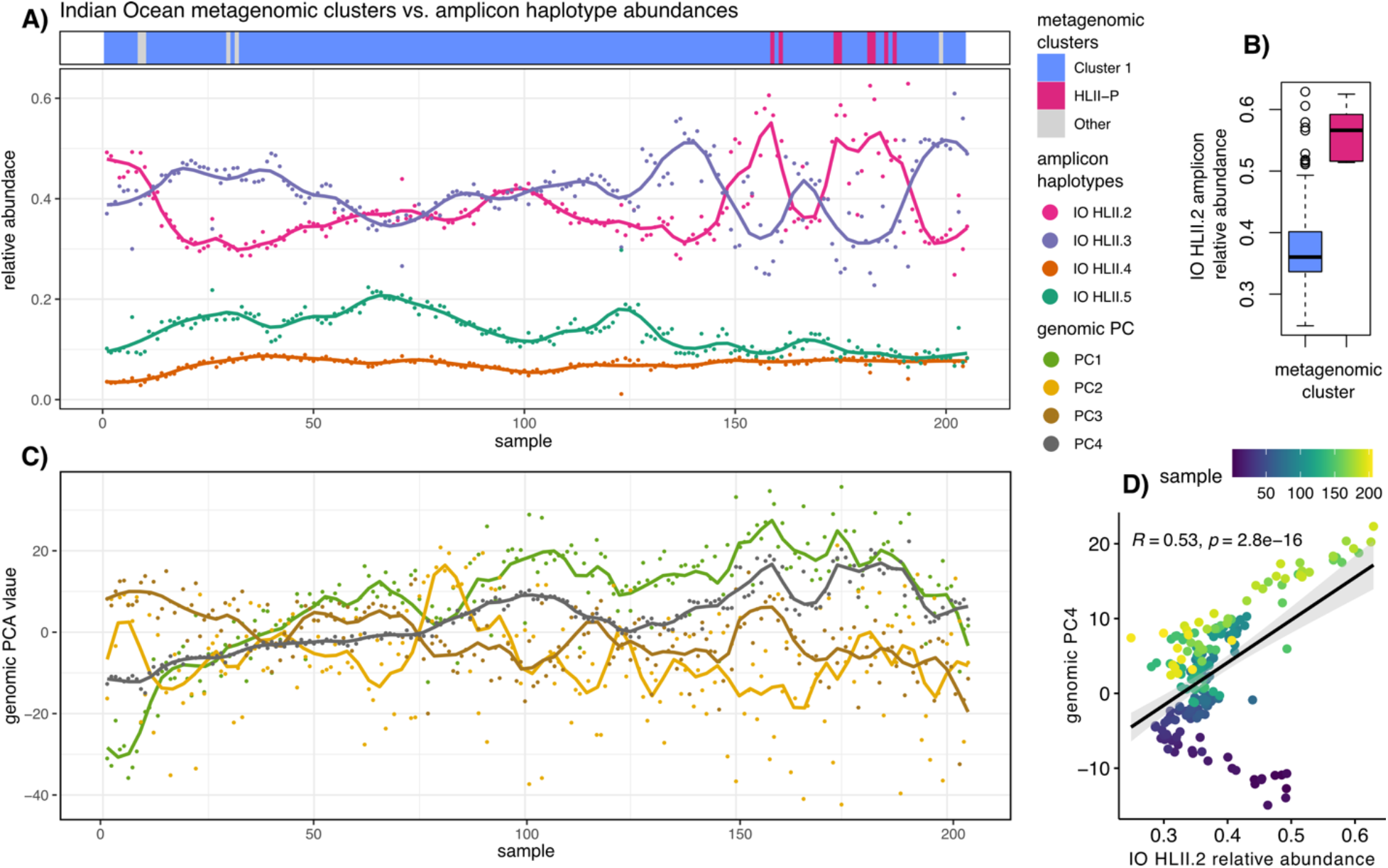
Comparison between amplicon and metagenomic phylogenetic diversity. (A) Metagenomically derived clusters compared to amplicon-based haplotype abundances (Larkin 2020). (B) Boxplot of IO HLII.2 amplicon sequence abundance grouped by metagenomic clusters. (C) Metagenomically derived genomic PCA values. (D) Linear comparison between IO HLII.2 relative abundance a metagenomically derived genomic PC4. Pearson correlation R value and p score are both show in the figure.

## Supplemental Tables

**Table S1:** Spearman correlation between genomic PCA analysis, nutrient gene abundances, and environmental factors.

|  | PC1 | PC2 | PC3 | PC4 |
| --- | --- | --- | --- | --- |
| **P genes** | p= 2.2e-16<br>rho= 0.3877267 | p= 0.0001964<br>rho= 0.147982 | p= 0.04191<br>rho= -0.0810904 | p= 2.2e-16<br>rho= 0.8771648 |
| **Fe genes** | p= 7.31e-06<br>rho= -0.1775951 | p= 5.944e-09<br>rho= -0.2291705 | p= 0.0003255<br>rho= -0.1427433 | p= 2.2e-16<br>rho= -0.7884059 |
| **N genes** | p= 5.638e-06<br>rho= -0.1800382 | p= 9.578e-08<br>rho= -0.2110631 | p= 2.2e-16<br>rho= 0.3912358 | p= 2.2e-16<br>rho= -0.4713667 |
| **Temp** | p= 2.2e-16<br>rho= 0.739164 | p= 0.0009222<br>rho= -0.134813 | p= 2.2e-16<br>rho= -0.3827844 | p= 2.2e-16<br>rho= 0.4637373 |

**Table S2:** Gene Content PCA (Genomic Diversity) Loadings of Nutrient Uptake and Metabolism Genes of *Prochlorococcus* HLII. PC rank is ordered based on the magnitude of the loading (absolute value).

| Gene | Function | Type | PC1 Rank | PC1 Loading | PC2 Rank | PC2 Loading | PC3 Rank | PC3 Loading | PC4 Rank | PC4 Loading |
| --- | --- | --- | --- | --- | --- | --- | --- | --- | --- | --- |
| *cirA* | Outer membrane receptor for ferrienterochelin and colicins | Fe | 1678 | 0.0106 | 2674 | 0.0017 | 2592 | -0.0021 | 55 | -0.0631 |
| *expD* | Biopolymer transport protein ExbD | Fe | 1617 | 0.0111 | 2659 | -0.0018 | 2579 | 0.0022 | 61 | -0.0613 |
| *febB* | ABC-type Fe3+-hydroxamate transport system, periplasmic component | Fe | 2162 | 0.0070 | 2542 | -0.0028 | 2397 | -0.0036 | 45 | -0.0646 |
| *fepB* | ABC-type cobalamin/Fe3+-siderophores transport system, ATPase component | Fe | 2303 | 0.0061 | 2319 | -0.0047 | 2429 | -0.0033 | 46 | -0.0645 |
| *fepC* | ABC-type Fe3+-siderophore transport system, permease component | Fe | 2246 | 0.0064 | 2259 | 0.0052 | 2046 | -0.0060 | 24 | -0.0676 |
| *fur* | Fe2+ or Zn2+ uptake regulation protein | Fe | 532 | 0.0273 | 127 | 0.0364 | 2172 | -0.0052 | 1531 | -0.0065 |
| *futB* | ABC-type Fe3+ transport system, permease component | Fe | 284 | 0.0320 | 63 | 0.0422 | 2828 | 0.0005 | 2562 | -0.0014 |
| *isiA* | Iron stress-induced chlorophyll-binding protein | Fe | 306 | 0.0315 | 819 | 0.0204 | 61 | -0.0426 | 1181 | -0.0087 |
| *isiB* | Flavodoxin | Fe | 351 | 0.0306 | 2837 | -0.0004 | 1908 | -0.0069 | 323 | -0.0287 |
| *tolQ* | Biopolymer transport protein ExbB/TolQ | Fe | 2231 | 0.0065 | 2541 | -0.0028 | 2742 | 0.0011 | 25 | -0.0672 |
| *tonB* | Lipoprotein-anchoring transpeptidase ErfK/SrfK | Fe | 2365 | 0.0056 | 2385 | -0.0042 | 2026 | -0.0061 | 68 | -0.0591 |
| *amt1* | Ammonia transporter | N | 595 | 0.0260 | 322 | 0.0298 | 199 | -0.0346 | 2134 | -0.0033 |
| *carA* | Carbamoylphosphate synthase small subunit | N | 174 | 0.0345 | 493 | 0.0260 | 1335 | 0.0124 | 2569 | -0.0013 |
| *cynA* | Cyanate transporter | N | 1435 | -0.0124 | 1597 | 0.0116 | 46 | 0.0447 | 264 | -0.0320 |
| *cynS* | Cyanate lyase | N | 1409 | -0.0126 | 1527 | 0.0123 | 40 | 0.0454 | 284 | -0.0311 |
| *dadA* | Glycine/D-amino acid oxidase (deaminating) | N | 477 | 0.0282 | 1840 | -0.0092 | 1189 | 0.0143 | 1689 | -0.0055 |
| *glnA* | Glutamine synthetase | N | 764 | 0.0224 | 31 | 0.0449 | 1146 | -0.0151 | 2423 | 0.0020 |
| *glnB* | Nitrogen regulatory protein PII | N | 262 | 0.0325 | 1065 | -0.0175 | 1583 | 0.0099 | 1411 | -0.0073 |
| *moaA* | Molybdopterin cofactor biosynthesis protein | N | 2744 | 0.0020 | 2768 | -0.0009 | 900 | 0.0189 | 331 | -0.0284 |
| *moaB* | Molybdopterin cofactor biosynthesis protein | N | 2727 | -0.0021 | 2181 | 0.0059 | 838 | 0.0199 | 353 | -0.0266 |
| *moaC* | Molybdopterin cofactor biosynthesis protein | N | 2813 | -0.0011 | 2008 | -0.0077 | 444 | 0.0279 | 274 | -0.0317 |
| *moaE* | Molybdopterin cofactor biosynthesis protein | N | 2575 | 0.0037 | 1290 | -0.0149 | 395 | 0.0294 | 341 | -0.0279 |
| *napA* | Nitrate/nitrite transporter | N | 2886 | 0.0000 | 2581 | -0.0025 | 861 | 0.0195 | 285 | -0.0310 |
| *narB* | Nitrate reductase | N | 2715 | 0.0022 | 2162 | -0.0062 | 692 | 0.0226 | 312 | -0.0290 |
| *nirA* | Nitrite reductase | N | 2851 | -0.0005 | 2811 | -0.0006 | 630 | 0.0238 | 304 | -0.0294 |
| *ntcA* | Nitrogen stress regulator | N | 909 | 0.0194 | 9 | 0.0494 | 1028 | -0.0168 | 2329 | 0.0025 |
| *pipX* | Nitrogen stress regulator | N | 253 | 0.0328 | 2564 | 0.0026 | 1021 | 0.0170 | 2084 | 0.0035 |
| *speA* | Arginine decarboxylase (spermidine biosynthesis) | N | 605 | 0.0258 | 2015 | 0.0077 | 894 | 0.0190 | 1023 | -0.0102 |
| *speB* | Arginase family enzyme | N | 2634 | -0.0031 | 1852 | 0.0091 | 387 | 0.0297 | 217 | -0.0363 |
| *ureA* | Urease gamma subunit | N | 243 | 0.0331 | 917 | -0.0192 | 1076 | 0.0161 | 2186 | -0.0031 |
| *ureB* | Urease beta subunit | N | 252 | 0.0328 | 1314 | 0.0145 | 2624 | -0.0019 | 1722 | -0.0053 |
| *ureC* | Urease alpha subunit | N | 589 | 0.0261 | 95 | 0.0383 | 708 | -0.0224 | 2798 | 0.0004 |
| *ureD* | Urease accessory protein UreH | N | 395 | 0.0297 | 641 | -0.0230 | 184 | 0.0350 | 1409 | -0.0073 |
| *ureE* | Urease accessory protein UreE | N | 547 | 0.0270 | 393 | -0.0280 | 280 | 0.0324 | 975 | -0.0107 |
| *ureF* | Urease accessory protein UreF | N | 398 | 0.0297 | 742 | -0.0216 | 324 | 0.0311 | 2157 | -0.0032 |
| ***ureG*** | Ni<sup>2+</sup>-binding GTPase involved in regulation of expression and maturation of urease and hydrogenase | N | 237 | 0.0332 | 871 | 0.0197 | 1644 | 0.0093 | 1287 | -0.0080 |
| ***urtA*** | ABC-type branched-chain amino acid transport system, periplasmic component | N | 292 | 0.0319 | 586 | 0.0241 | 583 | -0.0248 | 2763 | 0.0006 |
| ***acr3*** | Arsenite efflux pump ArsB, ACR3 family | P | 2565 | 0.0038 | 1891 | 0.0088 | 319 | 0.0312 | 51 | 0.0635 |
| ***arsR*** | Putative arsenate stress transcriptional regulator | P | 1135 | 0.0156 | 2009 | 0.0077 | 2262 | -0.0045 | 48 | 0.0643 |
| ***chrA*** | Transporter, ChrA family | P | 1727 | 0.0103 | 1933 | 0.0083 | 1285 | 0.0131 | 29 | 0.0669 |
| ***gap1*** | Glyceraldehyde-3-phosphate dehydrogenase | P | 1266 | 0.0140 | 1654 | 0.0109 | 1811 | -0.0078 | 42 | 0.0653 |
| ***mfs*** | Transporter, multifacilitator family | P | 1400 | 0.0127 | 1525 | 0.0123 | 2123 | -0.0054 | 36 | 0.0663 |
| ***phoA*** | Alkaline phosphatase (phoA) | P | 2393 | 0.0054 | 1528 | 0.0123 | 734 | 0.0218 | 31 | 0.0667 |
| ***phoB*** | P stress two-component response regulator | P | 1596 | 0.0113 | 2062 | 0.0072 | 1608 | 0.0096 | 28 | 0.0670 |
| ***phoE*** | Phosphate uptake outer membrane porin | P | 1193 | 0.0148 | 2135 | 0.0065 | 1903 | -0.0069 | 54 | 0.0632 |
| ***phoR*** | P stress two-component histidine kinase | P | 1545 | 0.0115 | 2268 | 0.0051 | 1383 | 0.0120 | 27 | 0.0670 |
| ***phoX*** | Alkaline phosphatase (phoX) | P | 1240 | 0.0142 | 2720 | -0.0014 | 2071 | -0.0058 | 90 | 0.0546 |
| ***PMM707*** | Hypothetical (upregulated under P stress) | P | 2426 | 0.0052 | 1625 | 0.0112 | 592 | 0.0246 | 37 | 0.0661 |
| ***PMM719*** | Hypothetical (upregulated under P stress) | P | 1462 | 0.0122 | 1506 | 0.0125 | 188 | 0.0348 | 351 | 0.0273 |
| ***PMM721*** | Hypothetical (upregulated under P stress) | P | 2561 | 0.0039 | 1636 | 0.0111 | 47 | 0.0446 | 95 | 0.0533 |
| ***PMM722*** | Hypothetical (upregulated under P stress) | P | 1078 | 0.0163 | 2754 | -0.0011 | 42 | 0.0451 | 178 | 0.0406 |
| ***ppA*** | Inorganic pyrophosphatase | P | 635 | 0.0252 | 30 | 0.0456 | 830 | -0.0201 | 2173 | -0.0031 |
| ***pstS*** | ABC-type phosphate transport system, periplasmic component | P | 1129 | 0.0156 | 727 | 0.0218 | 1239 | -0.0136 | 171 | 0.0421 |
| ***ptrA*** | P stress regulator | P | 2711 | 0.0023 | 1770 | 0.0098 | 189 | 0.0348 | 72 | 0.0582 |
| ***unkP1*** | Hypothetical (upregulated under P stress) | P | 1841 | 0.0095 | 1551 | 0.0121 | 1677 | 0.0089 | 39 | 0.0656 |
| ***unkP2*** | Hypothetical (upregulated under P stress) | P | 873 | 0.0202 | 1344 | 0.0141 | 399 | 0.0293 | 107 | 0.0520 |
| ***unkP3*** | Hypothetical (upregulated under P stress) | P | 2686 | 0.0026 | 1687 | 0.0106 | 351 | 0.0307 | 62 | 0.0610 |
| ***unkP4*** | Hypothetical (upregulated under P stress) | P | 2625 | -0.0032 | 1657 | 0.0108 | 91 | 0.0399 | 112 | 0.0516 |
| ***unkP5*** | Hypothetical (upregulated under P stress) | P | 1845 | 0.0094 | 2219 | 0.0055 | 678 | 0.0228 | 18 | 0.0692 |

**Table S3:** Reference Genome Database

| Genome | Accession | Species |
| --- | --- | --- |
| EQPAC1 | GCA_000759875.1 | *Prochlorococcus* |
| GP2 | GCA_000759885.1 | *Prochlorococcus* |
| HNLC1 | GCA_000218705.1 | *Prochlorococcus* |
| HNLC2 | GCA_000218745.1 | *Prochlorococcus* |
| MED4 | GCA_000011465.1 | *Prochlorococcus* |
| MIT0601 | GCA_000760175.1 | *Prochlorococcus* |
| MIT0602 | GCA_000760195.1 | *Prochlorococcus* |
| MIT0604 | GCA_000757845.1 | *Prochlorococcus* |
| MIT0701 | GCA_000760295.1 | *Prochlorococcus* |
| MIT0801 | GCA_000757865.1 | *Prochlorococcus* |
| MIT1312 | GCA_001632005.1 | *Prochlorococcus* |
| MIT1313 | GCA_001632065.1 | *Prochlorococcus* |
| MIT1318 | GCA_001632045.1 | *Prochlorococcus* |
| MIT1327 | GCA_001632125.1 | *Prochlorococcus* |
| MIT1342 | GCA_001632145.1 | *Prochlorococcus* |
| MIT9123 | GCA_000759935.1 | *Prochlorococcus* |
| MIT9201 | GCA_000759955.1 | *Prochlorococcus* |
| MIT9211 | GCA_000018585.1 | *Prochlorococcus* |
| MIT9215 | GCA_000018065.1 | *Prochlorococcus* |
| MIT9301 | GCA_000015965.1 | *Prochlorococcus* |
| MIT9302 | GCA_000759975.1 | *Prochlorococcus* |
| MIT9303 | GCA_000015705.1 | *Prochlorococcus* |
| MIT9312 | GCA_000012645.1 | *Prochlorococcus* |
| MIT9313 | GCA_000011485.1 | *Prochlorococcus* |
| MIT9314 | GCA_000760035.1 | *Prochlorococcus* |
| MIT9322 | GCA_000760075.1 | *Prochlorococcus* |
| MIT9401 | GCA_000760095.1 | *Prochlorococcus* |
| MIT9515 | GCA_000015665.1 | *Prochlorococcus* |
| NATL1A | GCA_000015685.1 | *Prochlorococcus* |
| NATL2A | GCA_000012465.1 | *Prochlorococcus* |
| PAC1 | GCA_000760235.1 | *Prochlorococcus* |
| RS50 | GCA_001989415.1 | *Prochlorococcus* |
| SB | GCA_000760115.1 | *Prochlorococcus* |
| SCGCAAA795_I06 | NA | *Prochlorococcus* |
| SCGCAAA795_I15 | NA | *Prochlorococcus* |
| SCGCAAA795_M23 | NA | *Prochlorococcus* |
| SS120 | GCA_000007925.1 | *Prochlorococcus* |
| UH18301 | SAMN00011132 | *Prochlorococcus* |
| XMU1401 | GCA_002812945.1 | *Prochlorococcus* |
| XMU1403 | GCA_003208065.1 | *Prochlorococcus* |
| XMU1408 | GCA_003208055.1 | *Prochlorococcus* |
| A9spades | NA | *Pelagibacter* |
| AAA240_E13 | SAMN02597172 | *Pelagibacter* |
| AAA288_E13 | SAMN02597171 | *Pelagibacter* |
| AAA288_G21 | SAMN02597281 | *Pelagibacter* |
| AAA288_N07 | SAMN02597280 | *Pelagibacter* |
| AAA298_D23 | GCA_000402655.1 | *Pelagibacter* |
| AG_337_G04 | ERS3879006 | *Pelagibacter* |
| AG_337_G06 | ERS3879008 | *Pelagibacter* |
| AG_426_I15 | ERS3880727 | *Pelagibacter* |
| AG_430_F16 | ERS3880946 | *Pelagibacter* |
| AG_430_I06 | ERS3880974 | *Pelagibacter* |
| B4spades | NA | *Pelagibacter* |
| F4spades | NA | *Pelagibacter* |
| HIMB058 | SAMN02440920 | *Pelagibacter* |
| HIMB083 | SAMN02597166 | *Pelagibacter* |
| HIMB114 | SAMN02436217 | *Pelagibacter* |
| HIMB122 | SRS843558 | *Pelagibacter* |
| HIMB1321 | GCA_900177485.1 | *Pelagibacter* |
| HIMB140 | ERS787856 | *Pelagibacter* |
| HIMB4 | NA | *Pelagibacter* |
| HIMB5 | SAMN00016662 | *Pelagibacter* |
| HIMB59 | SAMN00010387 | *Pelagibacter* |
| HTCC1002 | SAMN02436088 | *Pelagibacter* |
| HTCC1013 | SAMN02441456 | *Pelagibacter* |
| HTCC1016 | SAMN02256429 | *Pelagibacter* |
| HTCC1040 | SAMN02256395 | *Pelagibacter* |
| HTCC1062 | SAMN02603690 | *Pelagibacter* |
| HTCC7211 | SAMN02436224 | *Pelagibacter* |
| HTCC7214 | SAMN02841172 | *Pelagibacter* |
| HTCC7217 | SAMN02841150 | *Pelagibacter* |
| HTCC8051 | SAMN02440710 | *Pelagibacter* |
| HTCC9022 | SAMN02440781 | *Pelagibacter* |
| HTCC9565 | GCA_012932795.1 | *Pelagibacter* |
| IMCC1322 | GCA_000024465.1 | *Pelagibacter* |
| IMCC9063 | SAMN02603337 | *Pelagibacter* |
| PRT004 | NA | *Pelagibacter* |
| SAR86B | GCA_000252545.1 | *Pelagibacter* |
| BL107 | GCF_000153805 | *Synechococcus* |
| CB0101 | GCA_000179235.2 | *Synechococcus* |
| CB0205 | GCA_000179255.1 | *Synechococcus* |
| CC9311 | GCF_000014585 | *Synechococcus* |
| CC9605 | GCF_000012625 | *Synechococcus* |
| CC9616 | GCF_000515235 | *Synechococcus* |
| CC9902 | GCF_000012505 | *Synechococcus* |
| GEYO | GCF_900473955 | *Synechococcus* |
| GFB01 | GCA_001039265.1 | *Synechococcus* |
| KORDI_100 | GCF_000737535 | *Synechococcus* |
| KORDI_49 | GCF_000737575 | *Synechococcus* |
| KORDI_52 | GCF_000737595 | *Synechococcus* |
| MITS9508 | GCF_001632165 | *Synechococcus* |
| MITS9509 | GCF_001631935 | *Synechococcus* |
| N19 | GCF_900474045 | *Synechococcus* |
| N32 | GCF_900473895 | *Synechococcus* |
| NKBG042902 | GCA_000715475.1 | *Synechococcus* |
| PCC7335 | GCA_000155595.1 | *Synechococcus* |
| RCC307 | GCF_000063525 | *Synechococcus* |
| REDSEA_S02_B4 | GCA_001628325.1 | *Synechococcus* |
| RS9916 | GCF_000153825 | *Synechococcus* |
| RS9917 | GCF_000153065 | *Synechococcus* |
| UW105 | GCF_900473935 | *Synechococcus* |
| UW106 | GCF_900474015 | *Synechococcus* |
| UW140 | GCF_900474295 | *Synechococcus* |
| UW179A | GCF_900473965 | *Synechococcus* |
| UW179B | GCF_900474245 | *Synechococcus* |
| UW69 | GCF_900474185 | *Synechococcus* |
| **UW86** | **GCF_900474085** | *Synechococcus* |
| **WH5701** | **GCA_000153045.1** | *Synechococcus* |
| **WH7805** | **GCF_000153285** | *Synechococcus* |
| **WH8016** | **GCF_000230675** | *Synechococcus* |
| **WH8020** | **GCF_001040845** | *Synechococcus* |
| **WH8102** | **GCF_000195975** | *Synechococcus* |
| **WH8109** | **GCF_000161795** | *Synechococcus* |
| **Och114** | **GCA_000014045.1** | *Roseobacter* |

